# Distinct Spiking Patterns of Excitatory and Inhibitory Neurons and LFP Oscillations in Prefrontal Cortex during Sensory Discrimination

**DOI:** 10.1101/629659

**Authors:** Hua-an Tseng, Xue Han

**Affiliations:** Biomedical Engineer Department, Boston University, Boston, MA, 02215

## Abstract

Prefrontal cortex (PFC) spike activity and local field potential (LFP) oscillation dynamics are broadly linked to various aspects of behavior. PFC neurons can encode the identity of sensory stimuli and related behavioral outcome in a range of sensory discrimination tasks. However, it remains largely unclear how different neuron subtypes and related LFP oscillation features are modulated in mice during sensory discrimination. To understand how excitatory and inhibitory neurons in PFC are selectively engaged during sensory discrimination and how they relate to LFPs oscillations, we used tetrode devices to probe well isolated individual PFC neurons, and LFP oscillations, in mice performing a three-choice auditory discrimination task. We found that a majority of the PFC neurons, 78% of a total of 711 individual neurons, exhibited sensory evoked responses that are context and task-progression dependent. Using spike waveforms, we classified these responsive neurons into excitatory and inhibitory neurons, and found that both neuron subtypes were transiently modulated, with individual neurons’ responses peaking throughout the entire task duration. While the number of responsive excitatory neurons remain largely constant throughout the task, an increasing fraction of inhibitory neurons were gradually recruited as trial progressed. Further examination of the coherences between individual neurons and LFPs revealed that inhibitory neurons in general exhibit higher spike-field coherence with LFP oscillations than excitatory neurons, first at higher gamma frequencies at the beginning of the task, and then at theta frequencies during the task, and finally across theta, beta and gamma frequencies at task completion. Together, our results demonstrate that while PFC excitatory neurons are continuously engaged during sensory discrimination, PFC inhibitory neurons are preferentially engaged as task progresses and selectively coordinated with distinct LFP oscillations. These results demonstrate increasing involvement of inhibitory neurons in shaping the overall PFC network dynamics as sensory discrimination progressed towards completion.

## Introduction

The prefrontal cortex (PFC) is known to be critically involved in decision making, and damage to the PFC leads to deficits in various cognitive performance [1-7]. Goal-orientated decision making involves a number of cognitive aspects, including detecting sensory stimuli, applying learned rules, and executing an outcome. PFC activities, both individual neurons’ spiking patterns and population local field potential (LFP) oscillation dynamics, have been correlated with many aspects of the decision making process, such as attention, sensory processing, rule utilization, working memory, task progression tracking, and result anticipation. Recent studies using optogenetics to manipulate the activity of genetically defined cell types have showed that different PFC cell types are associated with distinct aspects of cognitive tasks [8-11]. In general, excitatory neurons participate in various aspects of a task, whereas different subtypes of inhibitory neurons seem to be preferentially recruited during different stages of a task. Calcium imaging of PFC neurons in a go/no-go task further revealed that excitatory neurons exhibit heterogeneous responses, while inhibitory neurons tend to be more correlated within their subtypes [12, 13] presumably due to gap junction coupling [14]. Parvalbumin-expressing (PV) neurons were shown to respond to various aspect of a task [12, 15], especially to reward [10, 11], whereas somatostatin-expressing (SST) neurons tend to be more selective and respond primarily to sensory stimuli and motor activity [10, 12].

LFP oscillations in PFC have been associated with a range of cognitive functions, and related to oscillations in other cortical and subcortical areas. For example, theta oscillations (∼5-10Hz) are closely linked to working memory [16], and are thought to coordinate long range connections between PFC and the hippocampus [17]. PFC Beta oscillations (∼15-30Hz), largely associated with status-quo and rule application [18], are often synchronized between PFC and other cortical areas. Higher frequency gamma oscillations (∼35-100Hz) are found to be mainly local within the PFC, and are thought to be primarily involved in working memory [19] and attention [20].

It has been suggested that LFP oscillations may organize neurons into functional ensembles [21, 22]. For example, coherence between the PFC spikes and the sensory cortex LFPs was increased during covert attention [20], and spike-field coherence within the sensory cortex was found to be correlated with behavioral performance [23]. Recent optogenetic experiments showed that abnormal activity of inhibitory neurons can disrupt gamma oscillations in PFC and lead to cognitive deficiency[24]. While much of our knowledge of the PFC has been obtained in humans, monkeys, and rats, much less is known about PFC function in mice, a model organism with advanced genetic tools that allow detailed examination of the functional significance of distinct neuron subtypes. Here, we recorded both spikes and LFPs simultaneously in the PFC while mice were performing a 3-choice auditory discrimination task in an open field. With tetrode devices, we identified well-isolated individual PFC neurons and distinguish excitatory neurons from inhibitory neurons using spike waveform features. We found that a large fraction of mouse PFC neurons and LFP oscillations were dynamic modulated during the sensory discrimination task, as observed in other animal models and humans. Inhibitory neurons were increasingly recruited towards task completion, suggesting that the inhibitory neural network is particularly engaged as sensory discrimination progressed. In addition, we found that inhibitory neurons overall showed higher coherence with LFPs compared to excitatory neurons, consistent with the idea that the activity of inhibitory neurons are more significantly coupled with LFP oscillations than excitatory neurons.

## Results

### A majority of individual PFC neurons exhibited task related spiking activities during a three-choice sensory discrimination task

To understand how distinct cell types are recruited during sensory discrimination, we designed a 3-choice auditory discrimination task, which required freely moving mice to associate a specific auditory stimulus with a predefined “reward” location (Figure 1A). Briefly, mice self-initiated each trial by stepping into the “initiation” location to trigger one of the three auditory cues (10 kHz sine wave, 25 click/second, and 100 click/second), which was presented throughout the trial. After trial initiation, mice were given 5 seconds to reach the “reward” location on the other end of the arena to receive a reward (correct trial). If mice reached the other two incorrect “reward” locations (incorrect trial) or failed to reach any “reward” location within 5 seconds (incomplete trial, excluded in this study), they were presented with a 5-second timeout, with bright light illuminating all “reward” locations. After training, all mice maintained a performance of >60% correct rate over the recording period (Figure 1B). Most mice were able to complete each trial within 3 seconds, with an average reaction time of 1.31±0.57 seconds across all completed trials in 6 mice (Figure 1C).

**Figure 1.**
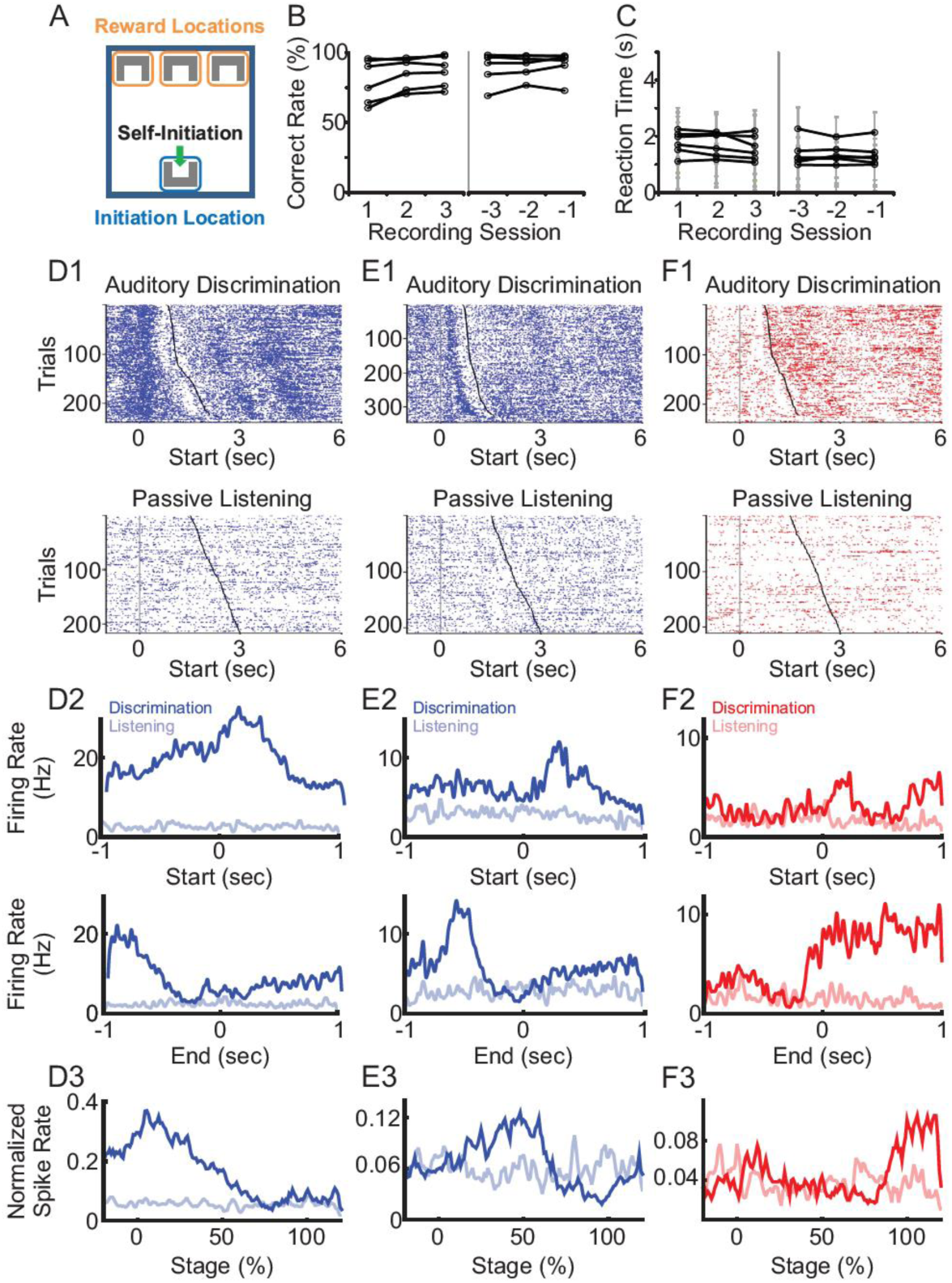
Excitatory and inhibitory neuron spiking during a three-choice auditory discrimination behavioral task. (A) During the auditory discrimination task, mice self-initiated each trial by triggering a motion detector at the “initiation” location. Upon trial-start, one of three auditory stimuli were presented throughout the trial. On the other end of the arena, there were three reward locations, each paired with a specific auditory stimulus. Mice were given 5 seconds to reach one of the three reward locations. If mice reached the correctly paired reward location, a reward was provided. If mice failed to reach the correct reward location, they experienced a 5 second timeout period, with a bright light illuminating the arena. (B) Behavioral performance of mice during the first three recording sessions (left) and the last three recording sessions (right). All mice had correct rates above random chance of 33% (N=6 mice). (C) The reaction time during the first three recording sessions (left) and the last three recording sessions (right). Each gray line represents mean±std of an individual mouse. (D) A representative excitatory neuron with elevated firing rate at trial start. Raster plots of spike activities from all trials were aligned to the trial start during the auditory discrimination tasks (D1 top) and during passive listening (D1 bottom). The average firing rates of the same neuron across all trials during the auditory discrimination tasks (dark line) and during passive listening (light line), aligned to the trial start (D2 top), and to the trial end (D2 bottom). Normalized firing rates throughout the entire trial duration of the discrimination task (D3 dark line) and of the passive listening block (D3 light line). (E) Similar to (D), but from a representative excitatory neuron with elevated firing rate in the middle of the task. (F) Similar to (D), but from a representative inhibitory neuron with elevated firing rate at the trial end.

We performed a total of 251 recording sessions in 6 mice, in PFC bilaterally, and identified 711 single neurons based on simultaneously recorded waveforms from four closely positioned tetrode wires (Fig. 1D). Among these 711 neurons, 552 (78%) showed significant changes in their firing rates during the task when compared to the inter-trial interval (ITI) (*p*<0.05, Wilcoxon rank sum test). To understand how different PFC neurons are selectively modulated during the sensory discrimination task, all subsequent analysis was performed on responsive neurons only.

We first examined the timing of PFC spiking relative to the stage of a trial. When aligned to trial start, some neurons showed an immediate increase in firing rate at trial start with the firing rate gradually decaying towards the end of the trial (Figure 1D), some exhibited delayed increase after trial start and mainly fired in the middle of the trial (Figure 1E), and some exhibited small increase at trial start but sharp rise towards the end of the trial (Figure 1F). Given the variable duration for a mouse to complete each trial, we normalized the firing rate to task duration. We found that individual neuron firing rates were dynamically modulated at different stages of the task. However, as a population, PFC neurons response covered the entire task duration, suggesting that the PFC neural network is engaged throughout the entire sensory discrimination task period (Figure 1D3, 1E3, and 1F3).

To rule out the possibility that PFC was solely driven by bottom up auditory stimuli, we designed a “passive listening” block, during which mice received the same auditory stimuli but without performing the discrimination task. Mice were placed in the same arena with a floor that covered the sensors at the “initiation” and “reward” locations. The “passive listening” block contained 200-250 trials. During each trial, one of the same three auditory stimuli was randomly presented for 1.5-3 seconds, followed by a 1.5-3 seconds long inter-trial interval (ITI). To compare the activity of the same neuron during discrimination task versus during passive listening, the “passive listening” block was performed either before or after the “discrimination task” block in the same recording session. Of the 422 task-modulated neurons that were also tested with the “passive listening” condition, only 12 (3%) neurons showed significant changes in firing rate (representative neurons shown in Figure 1D, E and F). Together, these results demonstrate that a large fraction of PFC neurons are selectively modulated during sensory discrimination. Even though individual PFC neurons are transiently activated during different phases of the task, the overall PFC neural network is engaged throughout the entire task period.

### Inhibitory neurons were preferentially modulated towards trial completion

To explore the difference in excitatory and inhibitory neural responses, we sorted the recorded individual neurons based on their spike waveforms. Of the 552 task-relevant neurons, the width of their spike waveforms followed a bi-normal distribution, with one peak centered at 0.45 ms, consistent with the general observation of putative excitatory neurons, and another peak centered at 0.25 ms, consistent with the general observation of putative inhibitory neurons (Figure 2A and B) [25-28]. Thus, based on spike width, we identified 441 excitatory neurons (80%) and 111 inhibitory neurons (20%), using 0.4 ms as a threshold. The recorded excitatory neurons exhibited a lower average firing rate comparing to the inhibitory neurons (excitatory: 1.66±1.84 Hz, inhibitory: 2.62±2.68 Hz; *p*<0.05, Wilcoxon Ranksum test).

**Figure 2.**
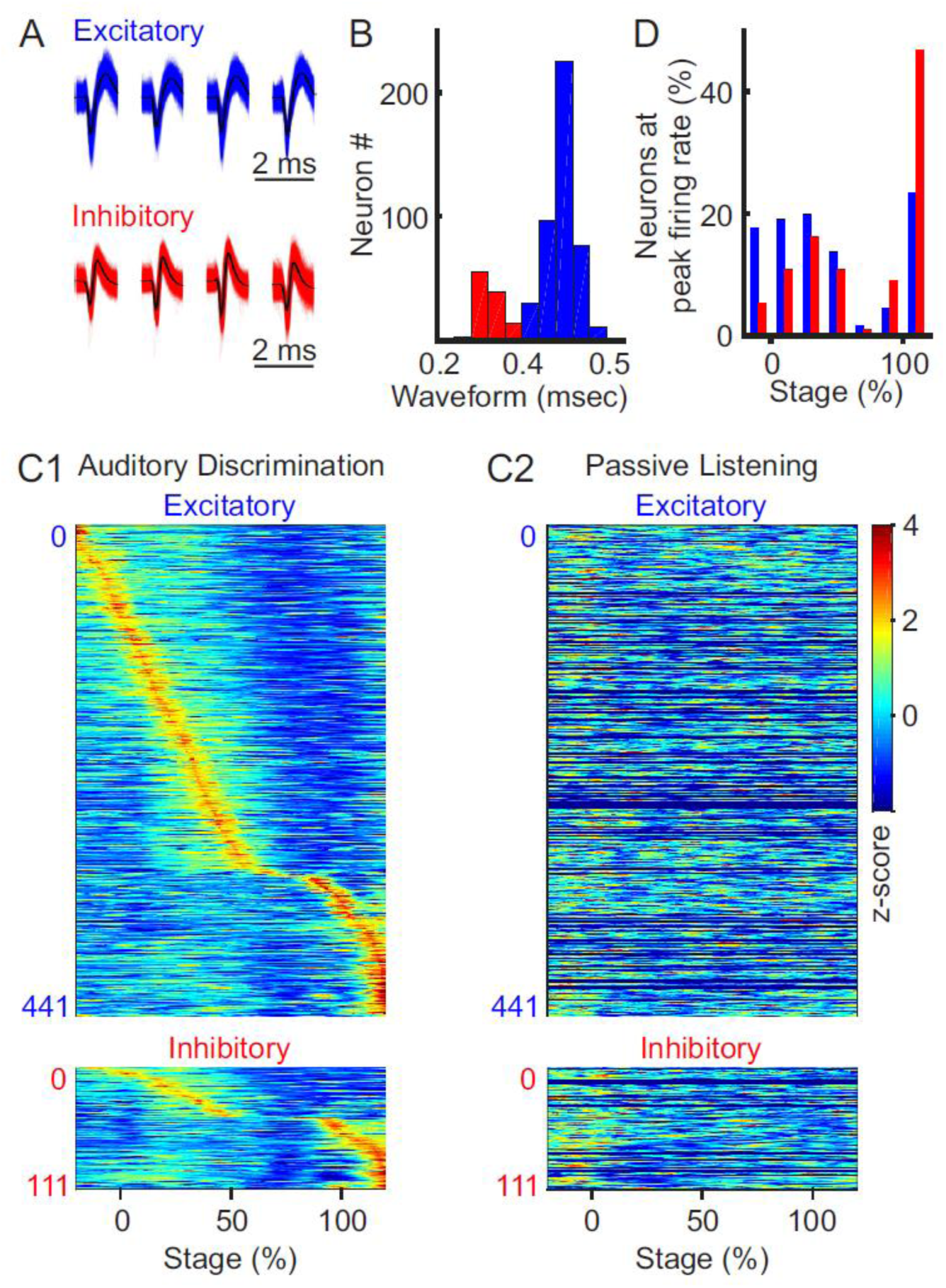
Context and stage-dependent modulation of spiking activity. (A) Representative waveforms of excitatory and inhibitory neurons recorded with tetrodes. (B) Binomial distribution of the width of spike waveforms, with a clear separation at 0.4 milliseconds between the two peaks, which was used as a threshold to identify excitatory (blue) and inhibitory (red) neurons. (C) Normalized population firing rate during the discrimination task (C1) and during passive listening (C2, sorted in the same order as in C1. 130 neurons were recorded only in auditory discrimination task without corresponding passive listening block were filled with dark blue). Neurons were grouped by type (excitatory and inhibitory) and sorted based on the timing of their peak firing rates. (D) Distribution of excitatory (blue bars) and inhibitory neurons (red bars), based on the timing of their peak firing rate during the task (*p*<0.01, *χ*^2^ test).

To examine how excitatory and inhibitory neurons are differentially modulated during the discrimination task, we sorted neurons according to the relative timing of their peak firing rates. We found that both excitatory and inhibitory neurons were transiently modulated throughout the task, with different neurons exhibit peak firing rate at different phases of the task (Figure 2C1). Excitatory neurons tended to be active throughout the entire task period, whereas increasing number of inhibitory neurons were recruited towards the end of the task (Figure 2D; *p*<0.01, χ^2^ test, between the distribution of excitatory and inhibitory neurons). On the other hand, these task responsive PFC neurons failed to produce any responses during the “passive listening” condition, confirming that PFC neurons exhibit task-specific modulation, rather than simply responding to bottom-up auditory stimuli alone (Figure 2C2). While PFC neurons are known to respond to auditory stimuli during passive conditions [29, 30], in our study, the three auditory stimuli at ∼70dB delivered over an ambient environment of ∼60dB were not sufficient to evoke significant passive responses in the PFC neurons. We further compared excitatory versus inhibitory neuron firing rates during correct versus incorrect trials, and found that both neuron subtypes showed clear sequential activity during the correct trials (Figure S1A1), but not during the incorrect trials (Figure S1A2), confirming that both excitatory and inhibitory neurons are modulated by task outcomes. Together, these results demonstrate that both excitatory and inhibitory neurons in the PFC encode task stage specific information that correlate with behavioral outcome. The fact that an increasing fraction of inhibitory neurons are recruited towards trial completion suggests that inhibitory network exhibits greater influence over the overall PFC dynamics towards the completion of the auditory discrimination as tested here.

### PFC neurons encode auditory stimulus identify, and inhibitory neuron populations exhibit increasing discrimination ability towards the completion of the task

While PFC neurons are broadly tuned to auditory stimuli, they are known to discriminate sensory stimuli and to categorize sensory inputs [5, 31, 32]. To examine whether mouse PFC neurons are selectively modulated by different auditory cues, we compared their responses to the three auditory cues used in the task (Figure 3A and B). We observed that PFC neurons exhibited highly heterogeneous responses to different cues. Some PFC neurons responded to only one auditory cue but not the other two (Figure 3A), whereas some responded to two cues but not the third one. This demonstrates that individual PFC neurons can encode cue identity with highly heterogeneous response profiles, highlighting that PFC networks could utilize the heterogeneity of individual neurons to expand the coding capacity of a large variety of cues among a population of individual neuron with various response amplitude and temporal kinetics. Consistent with the observation that PFC spiking responses are dynamic during the task, tone specific responses are often restricted to certain stages of the task.

**Figure 3.**
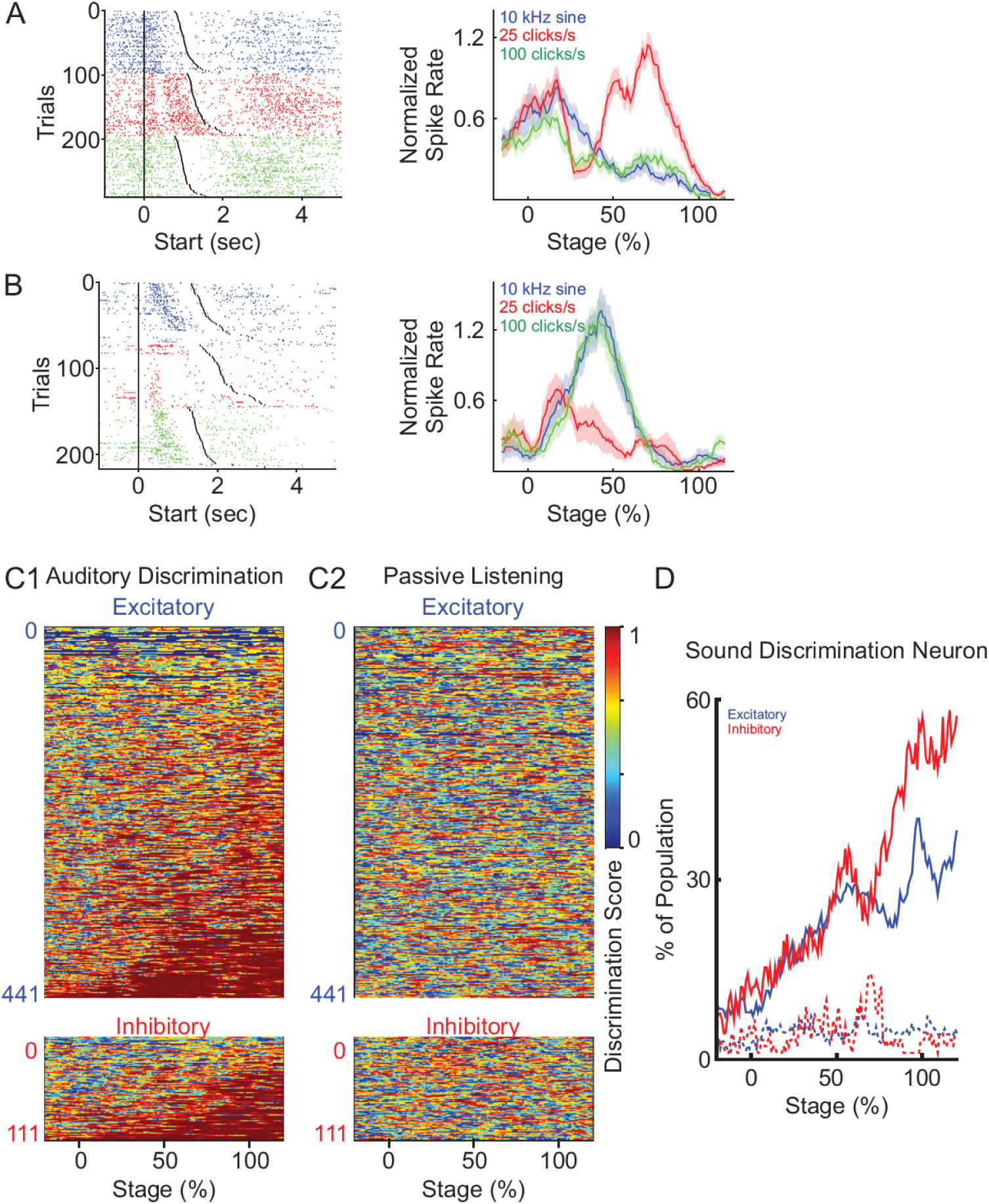
PFC neurons discriminate different auditory cues. (A) A representative neuron with increased response to the presentation of one auditory cue (25 click/sec), but not to the other two (10 kHz sine wave and 100 click/sec). *Left*: spike raster plot for trials with different auditory cues presentation (blue: 10 kHz sine wave; red: 25 click/sec; green: 100 click/sec), and sorted by trial durations. *Right*: normalized firing rate during each auditory stimulus across trials. (B) Another example neuron responded to two auditory cues (10 kHz sine wave and 100 click/sec), but not the third (25 click/sec). (C) Discriminatory ability of each neuron presented as discrimination score, defined as one minus the p-values calculated with one-way ANOVA between the firing rates upon the presentation of the three auditory cues at the different stages. Neurons were grouped as excitatory (top) and inhibitory neurons (bottom), and sorted by the total duration when they were discriminative. Discriminative responses of individual PFC neurons only occurred during the discrimination task (C1), but not during the passive-listening condition (C2, the neurons are sorted in the same order as C1). (D) The percentage of excitatory (blue) and inhibitory (red) neurons showing sound discrimination throughout trial stages.

To quantify the temporal auditory cue selectivity of each PFC neuron, we calculated the discrimination score, defined as 1 minus the *p*-value from One-way ANOVA test, between the firing rates upon the presentation of the three auditory cues at the different stages of the trial. A larger selectivity score indicates that the firing rates of a neuron exhibit greater difference in response to different sounds. We then plotted the selectivity scores of each neuron over the entire trial, and calculated the fraction of neurons that exhibit significant discriminatory activity during different task stages (Figure 3C1 and D). Overall, we found that an increasing number of excitatory and inhibitory neurons becomes sound discriminative as task progressed (Figure 3D). However, a significantly larger fraction of inhibitory neurons can distinguish tone identity toward the end of the task, compared to the excitatory neuron population (Figure 3D, excitatory: dark blue, inhibitory: dark red, χ ^2^ test, *p*<0.01). During control passive listening condition, these same PFC neurons failed to discriminate sound identity (Figure 3C2), and the percentage of modulated neurons stayed low and constant throughout the auditory stimuli presentation (Figure 3D, dot lines). Together, these results demonstrate that both excitatory and inhibitory PFC neurons spiking activity evolves to encode the identity of sensory stimuli during sensory discrimination, consistent with the idea that accumulating information converges onto PFC to facilitate the identification of sensory stimuli. In addition to recruiting a greater fraction of inhibitory neurons as sensory discrimination task progressed (Figure 2D), the proportion of inhibitory neurons exhibits sensory discrimination also increases more than excitatory neurons (Figure 3D), both of which results in a greater impact of inhibitory neural network over PFC overall network activity in facilitating sensory discrimination.

### Excitatory and inhibitory PFC neurons are differentially coordinated with PFC LFP oscillations at different frequency bands

PFC LFP oscillations have been broadly associated with many cognitive functions. We next examined LFP oscillation powers throughout auditory discrimination task, and observed that LFP power across all frequencies transiently decreased upon trial initiation, slightly recovered in the middle of the trial, and then sharply decreased again at trial end (Figures 4A). To quantify LFP oscillation changes, we compared the power at Theta (5-8 Hz), Beta (15-30 Hz), and Gamma (30-50 Hz) frequencies. Upon trial initiation, the reduction in oscillation powers was significant for all frequencies analyzed (Figure 4C; comparison between the averaged powers during the 500 ms time windows pre and post trial start: N=5 mice, paired t-test, *p*<0.05). At trial end, the decrease in oscillation power was also broad brand across all frequencies analyzed (Figure 4C; comparison between the averaged powers during the 500 ms time windows pre and post trial end: N=5 mice, paired t-test, *p*<0.05). Conversely, during passive listening, LFP powers seemed constant throughout the trial and progressed into the ITI (Figure 4B). However, similar analysis on the averaged power during pass listening revealed a small but significant increase across all frequencies at trial start (N=5 mice, paired t-test, *p*<0.05), in contrast to the reduction in oscillation powers at this stage during discrimination task (N=5 mice, paired t-test, *p*<0.05).

**Figure 4.**
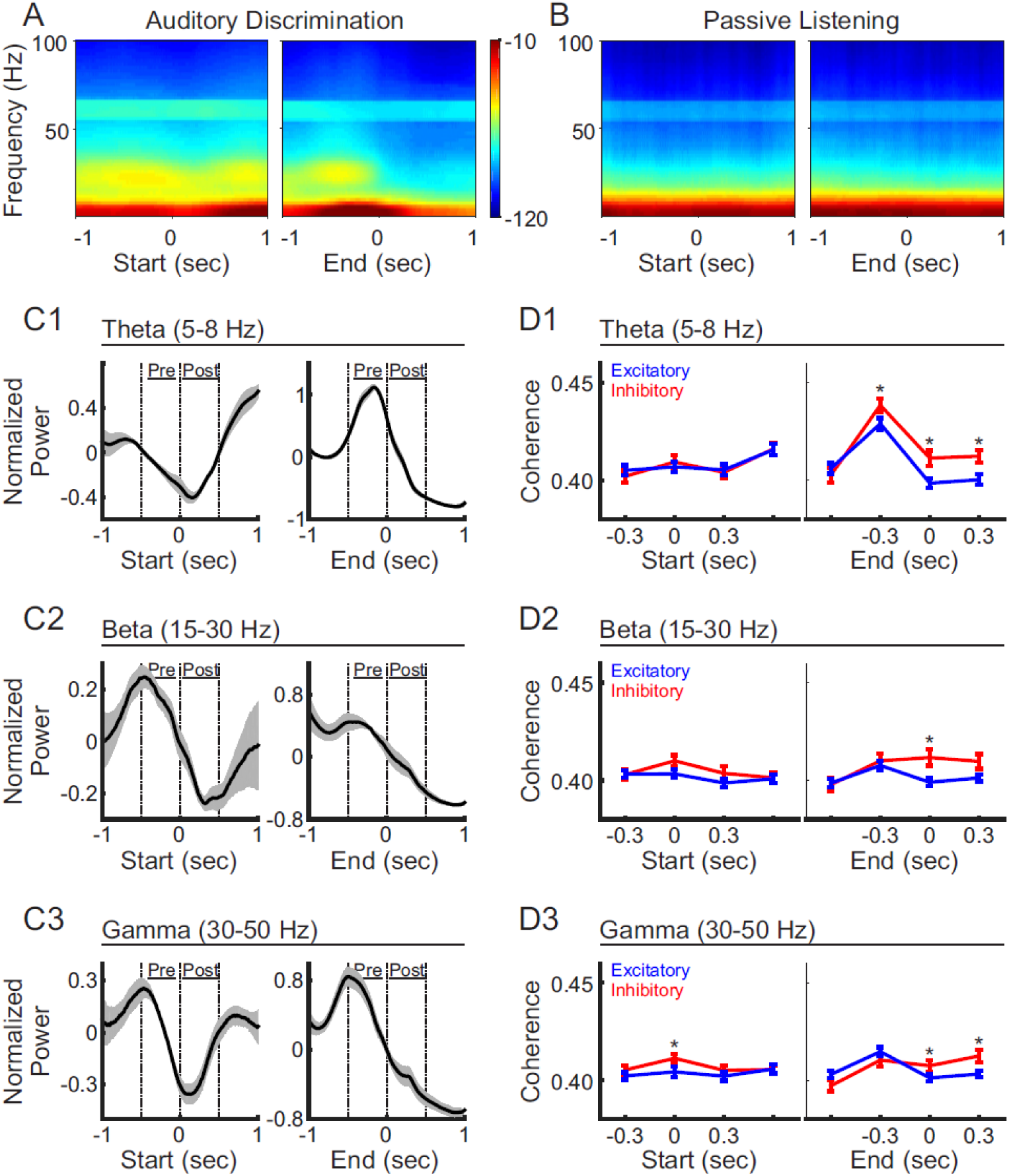
LFP oscillations and selective coherence with excitatory versus inhibitory neurons during auditory discrimination. (A) Average LFP power spectrum during the auditory discrimination from one example recording session, aligned at trial-start (left) and trial-end (right). (B) Average LFP power spectrum during passive listening from the same recording session as (A). (C) Normalized LFP oscillation powers at theta (C1, 5-8 Hz), beta (C2, 15-30 Hz), and gamma frequencies (C3, 30-50 Hz), aligned at trial-start (left), and at trial end (right), during auditory discrimination (black line). LFP powers were normalized as z-scores to the 2 second window analyzed. Shaded areas indicate standard error (N=5 animals). (D) Spike-field coherences of excitatory neurons (blue, mean±s.e.m) and inhibitory neurons (red, mean±s.e.m), aligned at trial-start (left), and at trial-end (right), during the discrimination task, at theta (D1), beta (D2) and gamma frequencies (D3), aligned to trial start (left) or to trial end (right). (Excitatory = 331 neurons, inhibitory= 91 neurons, t-test, *: *p*<0.05).

Even though LFP oscillations changes are broad band, inhibitory neurons are known to be related to specific oscillations [24]. To investigate the relationship between spiking activities of excitatory versus inhibitory neurons with LFPs, we calculated spike-field coherence between individual neurons and the LFPs recorded from the adjacent electrode within the ipsilateral hemisphere (Figure 4D). Interestingly, while LFP power is reduced across all frequencies at trial start, the reductions did not impact the coherence of excitatory and inhibitory neurons equally at all frequency bands. Inhibitory neurons exhibited significantly higher coherence only at gamma frequencies, but not at theta and beta frequencies (Figure 4D). This result is similar to the finding that inhibitory neurons are important for PFC gamma oscillations during rule shifting behavior [24], highlighting the general coupling of inhibitory neurons and PFC network dynamics during cognitive tasks. The fact that gamma frequency oscillation power is reduced at this stage, but yet inhibitory neurons are still more coherent with gamma frequencies, highlights that the PFC inhibitory network is preferentially engaged upon the initiation of the sensory discrimination task.

As trial progresses towards completion, spike-field coherence at theta frequency increased in both neuron types, with inhibitory neurons showing a higher coherence than excitatory neurons, which sustained beyond trial completion (Figure 4D1). At beta and gamma frequencies, the divergence of spike-field coherence between excitatory and inhibitory neurons only occurred after the trial end, where inhibitory neurons again showed stronger coherence than excitatory neurons (Figure 4D2 and 4D3). In summary, these results demonstrate that inhibitory neurons showed a stronger coherence with LFPs than excitatory neurons across multiple frequencies during the entire behavioral task, suggesting a more coordinated inhibitory neuron network that emerges with distinct LFP oscillations. It is possible that different inhibitory neuron subtypes are preferentially recruited as task progressed, which could account for the observation of the spike-field coupling with distinct frequency components as previously suggested [12-14].

### PFC LFP oscillation changes are outcome-dependent

To further understand the role of LFP oscillations in task performance, we investigated whether LFP oscillations were differently modulated by task outcome. When aligned to trial start, LFP oscillation power showed similar trends during the correct and incorrect trials (Figure 5A left vs 5B left, and 5C). We further quantified the dynamics of LFP power at trial start, by comparing the average power at theta, beta and gamma frequencies power during the 500 ms time windows before trial start versus the 500 ms window after trial start. We found that while the power is reduced on both correct and incorrect trials, the reduction is greater on correct trials than incorrect trials, across all frequencies (Figure 5D, N=5 mice, paired t-test, *: *p*<0.05).

**Figure 5.**
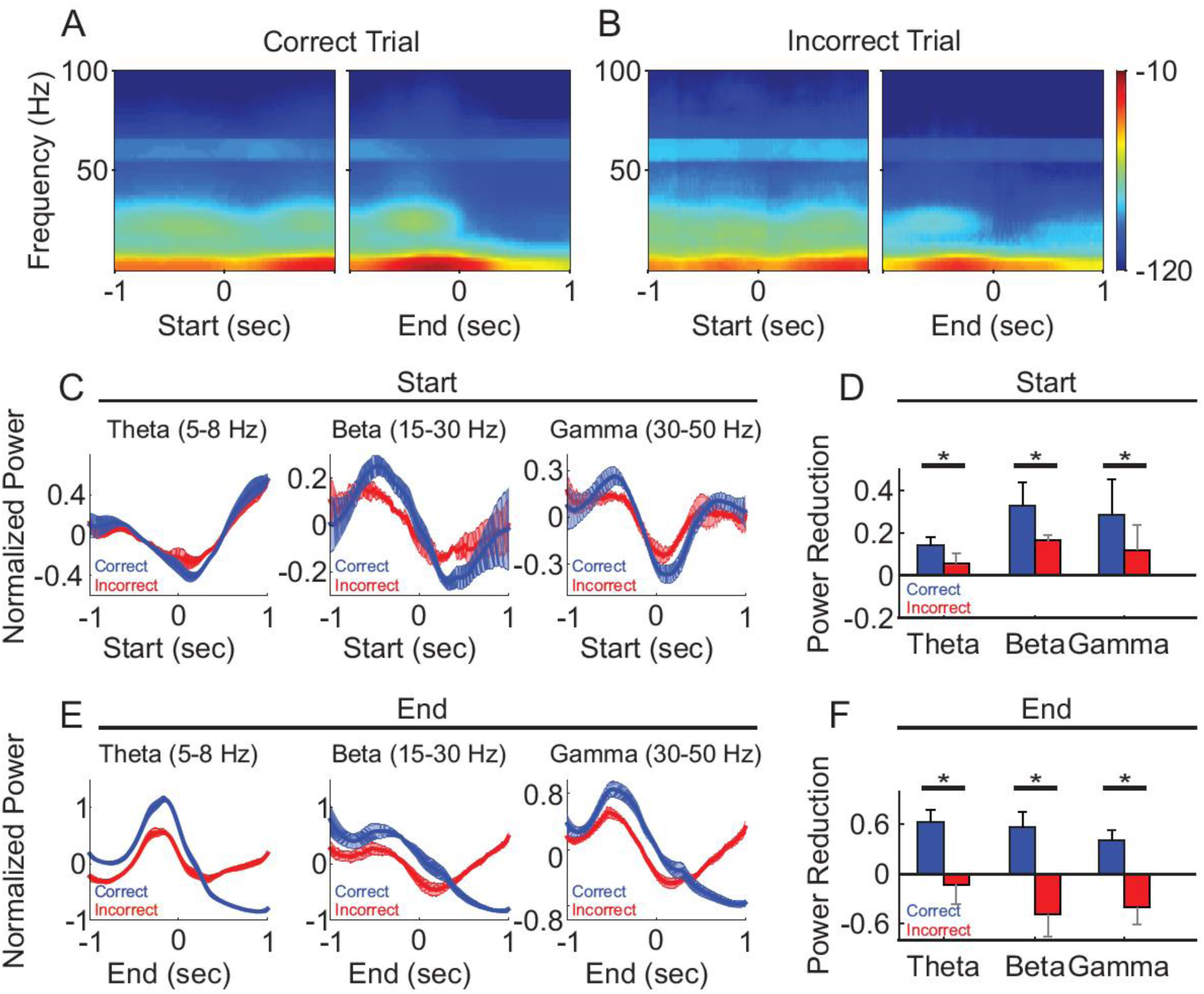
LFP oscillation power diverged between correct and incorrect trials. (A) Spectrogram of the correct trials from one representative recording session, aligned to trial-start (left) and to trial-end (right). (B) Spectrogram of the incorrect trials from the same recording session as (A), aligned to trial-start (left) and to trial-end (right). (C, E) Normalized population LFP oscillation powers of correct (black line) and incorrect (gray line) trials, aligned to trial-start (C) and to trial end (E). (D, F) The difference of LFP power, defined as the averaged z-scores of 500 ms windows before minus after at trial start (D), at trial end (F). (N=5, paired t-test, *: *p*<0.05).

Interestingly, at trial end, changes in oscillatory power diverged based on the trial outcome. On correct trials, oscillation powers continued to decrease and remained low over a prolonged period into the ITI, across all frequencies (Figure 5E, blue line). For incorrect trials, oscillation powers first decreased, similar to those in correct trials, but then rose across all frequency bands (Figure 5E, red line). As a result, the power changes at the trial end were bifurcated and significantly different, where the averaged powers showed reductions in correct trial, but exhibited an opposite relationship in incorrect trials (Figure 5F, N=5 mice, paired t-test, *: *p*<0.05). Together, these results demonstrate that LFP oscillation dynamics are linked to task performance, and the prolonged difference in LFP oscillations after each trial may serve as a feedback signal for the correct/reward and incorrect/timeout.

## Discussion

Understanding the different dynamics of excitatory versus inhibitory neural networks in PFC is of great interest in studying PFC involvement in cognitive functions. Here we designed a three-choice auditory discrimination task, and performed tetrode recordings from definitive single PFC neurons, in task performing mice. We distinguished well isolated excitatory neurons and inhibitory neurons based on spike waveforms, and found that inhibitory neurons were preferentially recruited at later stages, whereas excitatory neurons were active throughout the trial, but only during the discrimination task but not during passive listening. These results demonstrate that excitatory neurons and inhibitory neurons in PFC were recruited in a context-dependent manner at different stages of the auditory discrimination task. Not only an increasing faction of inhibitory neurons was recruited as task progressed, inhibitory neuron populations exhibited increasing discriminatory ability of the three auditory stimuli, and also selectively interacted with LFP oscillation at specific frequency bands and task stages. Together, these result demonstrate increasing impact of the inhibitory network in shaping overall PFC network dynamics during sensory discrimination.

PFC neurons exhibit diverse response profiles during cognitive tasks, from task progression [33], to different aspects or stages of a task [34]. PFC neurons increase their firing rates when anticipating task-relevant sensory stimulation [35], responding to sensory stimuli presentation [36, 37], maintaining working memory [38-41], predicting and/or anticipating reward [42, 43], or outcome [44-47], which can linger into the inter-trial interval [48]. We here show that mouse PFC neurons exhibit transient increases at specific stages of a 3-choice discrimination task, suggesting that different subgroups of PFC neurons can be sequentially recruited to process task-relevant information. The 3-choice auditory discrimination task allows us to examine more complex features of the PFC encoding ability. We found that while some neurons increased their firing rate specifically to only one auditory cue, others could be modulated by a group of cues. The ability of individual neuron responding to multiple cues could expand the PFC’s coding capacity, as each specific sensory information could be collectively represented from the combined selectivity of an ensemble of neurons.

Although mice were presented with the same auditory cues in both the “discrimination task” and the “passive-listening” blocks, PFC was only modulated during the “discrimination task” block, indicating that PFC neurons’ activities are context-dependent. Such context-dependent modulation has been shown when animals were exposed to different environments [49], or were performing tasks requiring different rules [50-54]. In our study, only a small fraction of PFC neurons (3% during passive listening vs 78% during discrimination task) was modulated during the “passive-listening” blocks, consistent with the idea of a behavioral state difference when animals were actively using sensory information in a task versus when passively presented with the sensory information. Such behavioral state difference may involve many neuromodulators, such as acetylcholine and dopamine that have been shown to mediate attentional states [46].

Different types of PFC neurons have been shown to exhibit distinct task-related responses [8, 9, 55]. Excitatory neurons increase their firing rates with less variability upon sensory stimulation [56], whereas inhibitory neurons are more correlated with the later stage of a task, such as reward [10]. Most recently, using optogenetics, Sparta et al. demonstrated that selective activation of PV neurons facilitated the extinction learning that involves the dissociation of the cue-reward relationships [11]. We found that an increasing proportion of neurons, both excitatory and inhibitory, were modulated by auditory cues as the task progressed, suggesting gradual recruitment of the PFC neurons during sensory discrimination, consistent with the idea that the PFC accumulates information about sensory identity and outcome decision [31]. Interestingly, we also found that inhibitory neurons showed higher spike-field coherence than excitatory neurons. Inhibitory neurons have been implicated in supporting LFP oscillations in the PFC [57, 58] and other brain areas [30, 59]. Our observation that a larger fraction of inhibitory neurons exhibited cue selectivity, together with the finding that inhibitory neurons possessed a stronger spike-field coherence, suggest that inhibitory neural network increasingly impact overall PFC network dynamics as discrimination task progressed.

While we found that LFP oscillatory powers were altered similarly across multiple frequency bands, the coherences between LFP oscillation and spike activity were neuron type specific. Inhibitory neurons showed stronger spike-field coherence than excitatory neurons with LFP oscillations at higher frequencies (gamma) at task start, which then switched to lower frequencies (theta) during the task. The fact that inhibitory neurons exhibited a higher degree of coherence across all frequency bands is suggestive of inhibitory neural networks in supporting PFC oscillation dynamics, which may play a crucial role in organizing cell assemblies within the PFC in a context-dependent manner as postulated for the general functional significance of LFP oscillations.

LFP oscillations at specific frequencies have been related to different aspects of behavioral tasks and states [13, 21, 60], and LFP synchrony within the PFC and between the PFC and other areas has been observed in many tasks [61]. In addition to single neuron responses in the PFC, we also observed wide spread changes in LFP oscillation patterns across all frequencies. At first glance, the oscillations across multiple frequency band seems to have similar power dynamics during the task, but a more detailed examination revealed that their coherences with the excitatory and inhibitory neurons differ, depending on the task stage and the frequency bands. Moreover, the dynamics of LFP power also reflected in task performance. As the LFP represents the collective activities from populations of neurons, a more phenomenal changes in oscillation could require a collaboration involving more PFC neurons and may be crucial in successfully performing the task.

## Materials and methods

### Three--choice auditory discrimination task

#### Behavior Arena

The behavior arena (12 inch × 12 inch) was constructed with black plastic walls and a mesh floor. The start location was located at one side of the arena (1 inch away from the wall), and three reward locations were at the opposite side (1 inch away from the wall) (Figure 1). Each location was equipped with an IR beam sensor and a white LED light on the floor. A speaker was located outside of the box, at the reward side, and delivered a 70 db auditory cue when the animal initiated the task. The room had a consistent background noise of approximately 60 db. LabVIEW (2012, National Instruments, Austin, TX) scripts were used to control the sensors and LED lights in the behavior arena via a NiDAQ board (NI USB-6259, National Instruments, Austin, TX) and recorded behavior events.

#### Behavioral training

All animal procedures were approved by the Boston University Institutional Animal Care and Use Committee. Female C57BL/6 mice (Taconic, Hudson, NY), were water-restricted during behavioral testing, and were closely monitored to ensure that they maintained at least 85% of their pre-experiment body weight. Adult female mice (2-3 months old at the start of the experiments) were trained to perform a 3-choice auditory discrimination task in following steps:

Step 1: Obtain water from reward port. At this step, only the middle reward port was accessible to the animal. It delivered a water reward whenever an animal reached the reward location and triggered the IR beam sensor in front of the water port.

Step 2: Detection of the first auditory cue. A white LED at the start location was illuminated to indicate trial initiation. Mice learned through trial and error to initiate a trial by reaching the start location, which triggered the IR beam sensor and the first auditory cue was presented. Mice were allowed to reach the reward location to obtain the reward without any time limit.

Step 3: Detection of the second auditory cue. After the animal learned initiating the task and responding to the first cue, we made the reward location associated with the second cue accessible, while blocking the reward locations for the other two cues. At this step, when animals initiated a trial, only the second cue was presented, and animals were required to reach the corresponding location for reward with no time limit.

Step 4: Two-choice auditory discrimination task. At this step, the reward locations for both the first and the second cues were accessible. When animals initiated the trial, one of two auditory cues would be presented randomly, and mice were required to reach the corresponding reward location for reward with no time limit. When animals reached the incorrect reward location, a 5 second timeout occurred, indicated by white LED lights around the reward locations.

Step 5: Detection of the third auditory cue: After the animal learned the 2-choice discrimination task, we repeated Step 3 to introduce the third auditory cue. At this step, only the third reward location was accessible and the other two were blocked. When an animal initiated the trial, only the third auditory cue was played, and the animal was required to reach the third reward location to complete the trial and receive the reward with no time limit.

Step 6: Three-choice auditory discrimination task. All three reward ports were accessible. When animals initiated the trial, one of the three auditory cues was presented, and mice were required to reach the corresponding reward location to obtain reward. At this stage, a reaction time limit of 5 seconds was introduced. Failure of reaching the reward location within 5 seconds would cause a timeout, indicated by white LED lights around the reward locations. To obtain a balanced number of trials with each cue, auditory cues were presented in a group of three, and within each group, each auditory cue was presented once in a random order.

During training, each mouse was trained 20 minutes per day. Once well trained, defined as performing over 60% correct rate per day over 3 consecutive training days and capable of completing a minimum of 100 trials per day, animals were provided free water access in their home cage for a week and then underwent tetrode implantation. After tetrode implantation and recovery from surgery, animals were briefly re-trained using procedures described in Steps 2-6 until their performance reached 60%, and then recordings were performed.

### Electrophysiology

Custom tetrode devices (16 channels) were assembled in house, which contained four tetrode bundles, two bundles targeting each hemisphere. The four tetrode bundles were designed to target the PFC bilaterally (AP: +1.8, ML: +0.2; AP: +2.2; ML +0.2; AP: 2.2; ML: −0.2; AP 2.2, ML: +0.2). A tetrode bundle was made by twisting together 4 tetrode wires (Sandvik-Kanthal, Ahmerst, NY), and adjusting the impedance to 0.5-1 MOhm with gold plating (SIFCO 5355, SIFCO ASC, Independence, Ohio). Tetrodes were implanted with the center positioned at the midline (AP: 2, ML: 0), so that the tetrode bundles targeted the PFC (AP: 2+/-0.2, ML: +/- 0.2). We advanced tetrode bundles gradually during the re-training period, so that the tip of the tetrode bundles reached the PFC at the recording stage (AP: 1.0 – 1.9). All recordings were performed in freely-moving mice. During recording, the tetrode device was connected to a commutator (ACO32, Tucker-Davis Technologies, Alachua, FL) to ensure free movement in the behavior arena. Data was acquired with a Plexon OmniPlex system (Plexon Inc, Dallas, TX). Spike waveforms and local field potentials were sampled at 40 kHz and at 1 kHz respectively. The Plexon OmniPlex system also received time stamps from the NiDAQ board to record timing of behavioral events.

Mice underwent one recording session per day. Each recording session constituted one “discrimination task” block and one “passive listening” block in random order. Animals could move freely in the behavioral arena during the entire recording session. The “discrimination task” block lasted about 20 minutes, and animals were allowed to initiate the task as many times as they desired. In general, mice performed 100-200 trials within 20 minutes. During the “passive listening” block, mice were placed in the same behavioral arena with a plastic floor positioned above all IR beam sensors and LEDs, so that mice had no access to any sensors or water ports. The same three auditory cues (∼70db) used in the “discrimination task” were played in pseudo-random order for 200-250 trials. The durations of the auditory cues were randomized from 1.5 to 3 seconds with random 1.5 - 3 seconds inter-trial intervals.

### Data analysis

Spike: Spikes were sorted with Offline Sorter (Plexon Inc, Dallas, TX) and then imported into Matlab (2014, MathWorks, Natick, MA) for further analysis. Spike width was defined as the duration between the valley and the peak of a spike waveform. Due to the difference in lengths of each trial, to calculate the firing rate throughout trial progression, we first normalized each trial based on its duration, so that trial start and trial end were aligned at 0% and at 100% of trial progression, respectively. We then calculated the firing rate from −20% to +120% of trial progression using a 1% moving window, and smoothed the results by averaging each data point with its two adjacent data points. When presenting the population data, we further normalized the firing rate between −20% to 120% of trial progression by calculating the z-scores for each neuron with its own mean and standard deviation.

LFP: LFPs were imported into Matlab with the Matlab custom script provided by Plexon, and then analyzed with the Chronux toolbox (chronux.org). The power spectrogram of each LFP trace was calculated with mtspecgramc function (moving window size: 500 ms, moving window step: 5ms, tapers: [3 5]) in Chronux.

In a few occasions, animal movement caused artifacts in our tetrode recording, such as when the tetrode devices bumped the arena walls, which create large voltage deflections in our recordings. To eliminate the impact of such movement artifact in our analysis, we identified these artifact periods as having the 5% maximum amplitude, either positive or negative, of the whole recording session. Trials that contained these artifact periods were excluded from all analysis involving LFPs.

The spectrogram examples were log-normalized (10*log(power/max)), with the maximum power of the examined time window (2 second, centered at either trial start or end). To compare the powers at specific frequency bands across mice, the powers within the examined time window (2 second, centered at either trial start or end) of each trial were first normalized by converting to their z-scores and then averaged within a given frequency band to obtain the power of each trial. The powers of all trials from the same animal were then averaged as the representative power of each animal. In our analysis, we examined three frequency bands, and the ranges of each frequency bands were defined as follows: theta (5-8 Hz), beta (15-30 Hz), and gamma (30-50 Hz).

Spike-field coherence was calculated with cohgramcpt function (moving window size: 500 ms, moving window step: 5ms, tapers: [3 5]) in Chronux. For each trial, we first calculated the spike-field coherence for each neuron with LFPs at a specific frequency range, and then averaged across trials.

### Statistical testing

For spike rate modulation, we used one-way ANOVA to compare the firing rates at the same trial progression period for different auditory cues. For LFP power spectrum, we used a paired t-test to compare the power during the 500 ms before and after either trial start or trial end. For spike-field coherence, we used a non-paired t-test to compare the coherence at the same trial progression period between the “discrimination task” blocks and “passive listening” blocks. All analyses were performed in Matlab. Details for each test are presented throughout the Results section.

## Data Availability

The datasets generated during and/or analyzed during the current study are available from the corresponding author on reasonable request.

## Acknowledgements

We thank members of Han lab for suggestions on the manuscript. This manuscript has been released as a pre-print at bioRxiv [62].

## Author Contributions

H.T and X.H. designed all experiments and wrote the paper. H.T. performed experiments and data analysis. X.H. supervised the project.

## Financial Disclosure

X.H. acknowledges funding from NIH 1R34NS111742, NSF 1835270 and 1848029, and Boston University Biomedical Engineering Department.

## Competing Interests

The author(s) declare no competing interests.

## Figures

**Figure S1.**
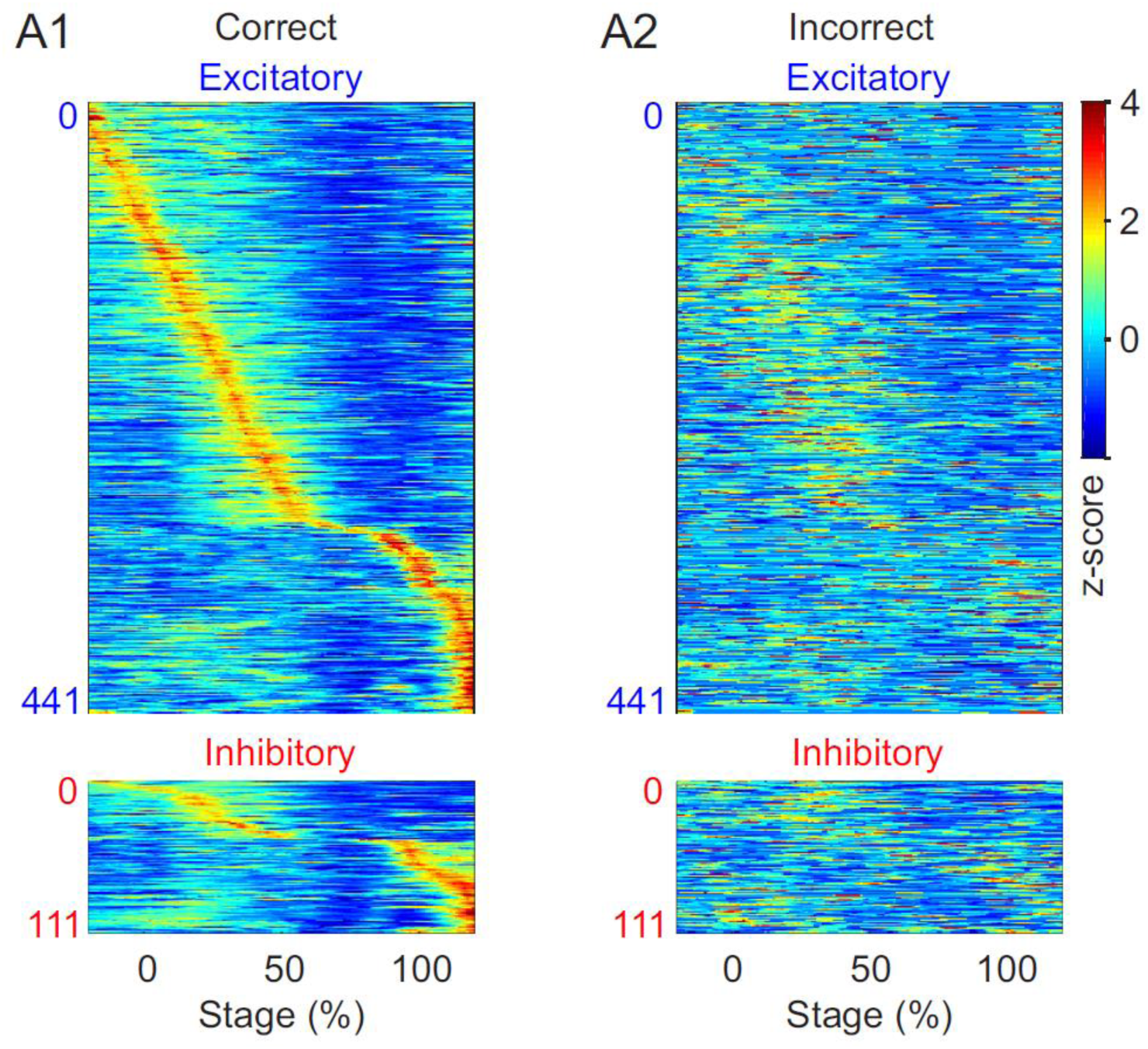
PFC spiking activity during correct versus incorrect trials. (Related to Figure 2) Normalized population firing rates of excitatory (blue) and inhibitory (red) neurons during correct trials (A), and incorrect trials (B).

## Notes

### Competing Interest Statement

The authors have declared no competing interest.

